# Aerosolization of *Mycobacterium tuberculosis* by tidal breathing

**DOI:** 10.1101/2021.10.17.464541

**Authors:** Ryan Dinkele, Sophia Gessner, Andrea McKerry, Bryan Leonard, Juane Leukes, Ronnett Seldon, Digby F. Warner, Robin Wood

## Abstract

**Rationale:** Interrupting tuberculosis (TB) transmission requires an improved understanding of how – and when – the causative organism, *Mycobacterium tuberculosis* (*Mtb*), is aerosolized. Although Cough is commonly assumed to be the dominant source of *Mtb* aerosols, recent evidence of Cough-independent *Mtb* release implies the contribution of alternative mechanisms.

**Objective:** To compare the aerosolization of *Mtb* and particulate matter from GeneXpert-positive patients during three separate respiratory manoeuvres: Tidal Breathing (TiBr), Forced Vital Capacity (FVC), and Cough.

**Methodology:** Bioaerosol sampling and *Mtb* detection were combined with real-time assessments of CO_2_ production and particle counts from 39 confirmed TB patients.

**Measurements and Main Results:** TiBr and FVC produced comparable numbers of particles, with Cough producing >4-fold more. For all manoeuvres, the proportions of particles detected across size categories from 0.5 – 5 μm were similar, with minor differences observed only in particles between 1.5 – 2 μm (p = 0.014) and >5 μm (p = 0.020). Viable *Mtb* bacilli were detected in 66%, 70%, and 65% of TiBr, FVC, and Cough samples, respectively. Notably, while Cough produced 3-fold more *Mtb* than TiBr, the relative infrequency of coughing compared to breathing implies that TiBr likely contributes >90% of the daily aerosolised *Mtb* across a range of Cough frequencies.

**Conclusions:** Our results suggest that, while Cough increases particle aerosolization compared to TiBr, this is not associated with increased *Mtb* aerosolization. Instead, TiBr produces more *Mtb* per particle than Cough. Assuming the number of viable *Mtb* organisms detected provides a proxy measure of patient infectiousness, these observations imply a significant contribution of TiBr to TB transmission.

## Introduction

Chronic Cough is a hallmark symptom of tuberculosis (TB), an airborne infectious disease which is caused by *Mycobacterium tuberculosis* (*Mtb*) and is associated with high global mortality and morbidity (1). Given its importance in TB diagnosis, Cough has unsurprisingly been central to TB transmission research (2). There are multiple lines of evidence, however, which suggest that the focus on Cough risks ignoring other important contributing mechanisms, undermining the implementation of new approaches to reducing TB transmission – especially in TB endemic settings. For example, a recent national TB prevalence survey in South Africa (consistently among the WHO’s annual list of high TB burden countries) reported that nearly 60% of individuals with bacteriologically confirmed pulmonary TB were asymptomatic (3). Similarly, a pioneering face-mask sampling study noted no association between Cough frequency and the detection of *Mtb* organisms (4). When considered with modelling data which estimate 1.7 billion latent *Mtb* infections globally (5), these observations imply that most *Mtb* infections are not associated with Cough as pathognomonic symptom – even when *Mtb* bacilli are present at detectable levels (3). Given the low proportion of infections leading to symptomatic disease and the scale of the TB epidemic, it is conceivable that *Mtb* transmission from symptomatic individuals, and therefore Cough, is highly efficient. However, considering the challenges in identifying TB transmitters (6) and the lack of association between *Mtb* detection and Cough frequency (4), an alternative must be considered: is TB transmission primarily accomplished by Cough-independent means?

We have developed a platform combining non-invasive bioaerosol capture technology and fluorescence microscopy to enumerate viable *Mtb* released by confirmed TB patients (7–9). Using this platform, we detected *Mtb* bacilli in the absence of (induced) Cough. More recently, we compared deep exhalations to Cough and found no difference in *Mtb* aerosolization (10). However, these observations were not conclusive given that we did not directly investigate the propensity for respiratory aerosol production in each manoeuvre. Therefore, we aimed in the current work to investigate the potential for *Mtb* release via Tidal Breathing (TiBr), Forced Vital Capacity (FVC) and Cough. As detailed below, our observations suggest that release of *Mtb* aerosols via TiBr might constitute an important contributor to ongoing transmission in both active TB disease and asymptomatic *Mtb* infection.

## Methods and materials

### Participant recruitment

Participants over 13 years of age returning a GeneXpert-positive sputum result were recruited from March 2020 to June 2021 at primary healthcare facilities in Ocean View and Masiphumelele, peri-urban townships in Cape Town, South Africa. Recruitment and sampling occurred prior to the initiation of standard anti-TB chemotherapy. Ethical approval was obtained from the Human Research Ethics Committee of the University of Cape Town (HREC 529/2019).

### Sample collection

Bioaerosols from three respiratory manoeuvres, Forced Vital Capacity (FVC), Tidal Breathing (TiBr), and Cough, were captured in liquid cyclone collectors utilising a direct sampling strategy (Figure 1A) (10). A unidirectional airflow forced exhaled air via a CO_2_ monitor and a high-flow cyclone collector at a maximum flow rate of 300 L/minute, trapping particulate matter in the collection medium (sterilised phosphate-buffered saline, PBS). During TiBr sampling, each participant placed their head within the elliptical cone and breathed normally for five minutes. Bioaerosols were captured within the cyclone collector at 200 L/minute, while 100 L/minute of exhaled air was diverted to an aerodynamic particle sizer (APS). During FVC and Cough sampling, each participant performed 15 manoeuvres by placing their head in the elliptical cone every 15 seconds. Bioaerosols were captured within the cyclone collector at 300 L/minute for the first and last five manoeuvres. During the middle five manoeuvres, 100 L/minute of exhaled air was diverted via the APS. New cyclone collectors were attached after each sample allowing for the independent enumeration of *Mtb* utilising our previously described concentration and visualisation pipeline (8).

**Figure 1.**
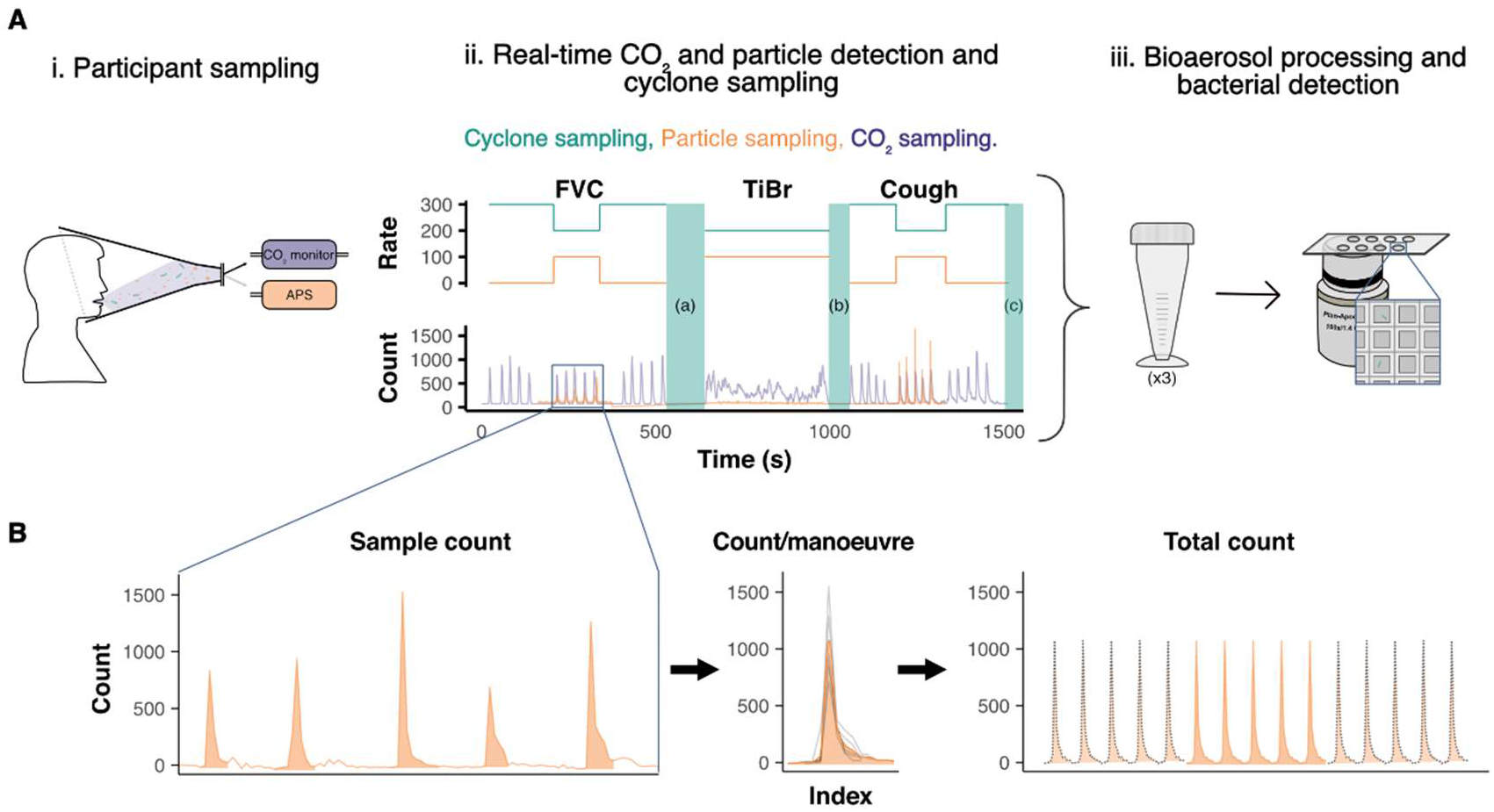
Participant sampling strategy. (**A**) i) GeneXpert-positive participants were recruited from TB clinics in Masiphumelele and Ocean View. Bioaerosol samples were collected in the RASC from three respiratory manoeuvres: Forced vital capacity (FVC), tidal breathing (TiBr) and coughing (Cough). ii) During sampling, real-time data were collected for CO_2_ concentration (purple) and particle production (orange). Measuring particles required diverting one-third (100 L/min) of the exhaled air into an aerodynamic particle sizer (APS). Particles were only counted for one-third of FVC and Cough sampling, and for the full duration of TiBr sampling. iii) Three independent liquid cyclone collectors were attached for each manoeuvre, which were microscopically scanned for *Mycobacterium tuberculosis* (a, b and, c within the green columns, green lines indicate sample collection flow rate). (**B**) Graphical representation of the variables; sample count, count/manoeuvre and total count. Sample count is the total count (volume) of particles sampled. Count(volume)/manoeuvre is the average count (volume) of particles per manoeuvre during particle sampling. Total count (volume) is the estimated total number of particles per sample, calculated by multiplying the average count per manoeuvre by the total number of manoeuvres during sampling (counted utilising CO_2_ data).

### Staining and visualisation of bioaerosol samples

Bioaerosols were stained with DMN-trehalose (DMN-tre) (Olilux Biosciences Inc.) and visualised as previously outlined (8). Briefly, liquid-captured bioaerosols (5 – 10 mL) were centrifuged for 10 minutes at 3000 × *g* (Allegra X-15R, Beckman Coulter) and resuspended in 200 μL of Middlebrook 7H9 medium supplemented with 100 μM DMN-tre. Staining was done overnight, after which samples were concentrated at 13,000 × *g* for five minutes and resuspended in 20 μL filtered PBS. Stained samples were loaded on nanowell-arrayed microscope slides and viewed on a Zeiss Axio Observer 7 with widefield illumination from a 475 nm LED and a Zeiss 38 HE filter set. A 100× plan-apochromatic phase 3 oil immersion objective (NA = 1.4) was used.

### Statistical methods

Each participant performed three respiratory manoeuvres violating the assumption of independence. Therefore, to account for this correlation within the data, various Linear Mixed Models were used as the incorporation of the random effect enabled the average differences between manoeuvres to be determined while accounting for variation between participants. For continuous outcomes, a log_10_-transformation was performed and linearity, normality of residuals, and homoskedasticity assessed. For binary outcomes, logistic regression was performed with sample type as the fixed effect, and variation in slope (random effects) accounted for by participant. For count data, negative binomial regression was applied. Detailed statistical and data wrangling methods can be found in the online supplement. Data were analysed in R studio (11) with R version 4.0.3 (12).

## Results

### Detection and quantification of particle release during different respiratory manoeuvres

Direct bioaerosol sampling was performed on 39 participants with corresponding CO_2_ and particle data obtained for 32 and 33 participants, respectively; this ensured a final sample size of 32 (Figure E1). FVC and Cough samples were excluded if fewer than two peaks in particle counts were detected above background. Owing to variations in sampling duration, samples were assessed in three ways (Figure 1B): the total number or volume of particles collected, the average number or volume of particles produced per manoeuvre, and the estimated total number or volume of particles produced.

During the particle sampling window, similar numbers (Figure 2A) and volumes (Figure E2A) of particles were collected for TiBr and Cough, with FVC producing significantly fewer particles than TiBr. However, after averaging the number of particles per manoeuvre, it was clear that TiBr produced a significantly lower number (Figure 2B) and volume (Figure E2B) of particles compared to either FVC or Cough. Interestingly, when considering the total number of manoeuvres performed, TiBr and FVC produced comparable numbers (Figure 2C) and volumes (Figure E2C) of particles, with Cough producing more than 4-fold more particles than TiBr. Inter-participant variability contributed approximately 12% of the total variation when examining particle numbers per manoeuvre (Figure 2B), and approximately 16% of the variation when examining particle volumes per manoeuvre (Figure E2B). Together, these data suggest that variation in particle production between the three respiratory manoeuvres contributed to variation in the overall volume of bioaerosol collected after 5 minutes of sampling; however, no manoeuvre was significantly under-sampled.

**Figure 2.**
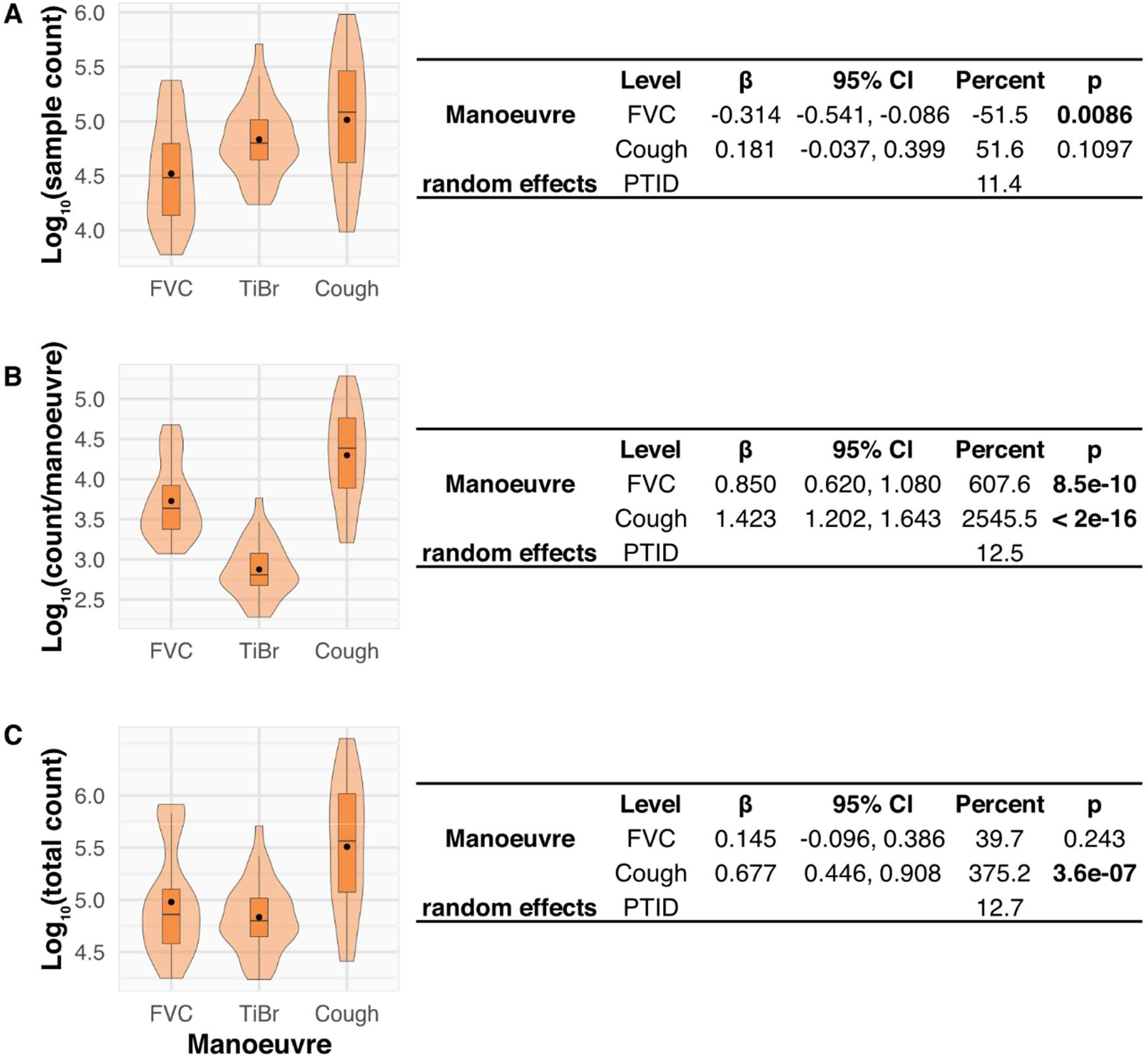
Variation in particle production by FVC, TiBr and Cough. A comparison of the (**A**) sample count, (**B**) count/manoeuvre and (**C**) total count of particles during sampling. The adjacent tables contain the results of univariate linear mixed models for each. The beta-coefficient (β) and 95% confidence interval (CI) are presented with percentage change relative to TiBr (Percent). The random effects results indicate the degree of variation (in %) between participants.

### Size stratification of particles enables more specific comparisons of respiratory manoeuvres

Particles of various sizes are aerosolised and may differ between respiratory manoeuvres. The APS binned particles into size categories (Figure 3A); therefore, we examined the effect of each manoeuvre on the distributions of particles across categories. The average number (Figure 3B) and volume (Figure E3A) of particles per manoeuvre were stratified by size category and sample type, with the individual data from each participant overlayed. Two features were apparent: firstly, there was a consistent distribution in average count per size category across all three manoeuvres. Moreover, this trend was recapitulated when the proportion of each size category relative to the total particle count per manoeuvre was compared for each size category (Figure 3C). Only minor variations were detected, with levels of significance reached solely for C.3 (1.5 – 2 μm) and C.5 (>5 μm). This suggested that factors leading to increased bioaerosol generation did not affect the distribution of size categories aerosolised over the size range measured. The second striking observation was the per patient consistency in the relationship between the different size categories. It was also apparent that size category was not a confounder of the relationship between the manoeuvre type and particle count per manoeuvre (Figure E3B), as the average differences between TiBr and FVC and TiBr and closely recapitulated those observed in the previous model (Figure 2B). In combination, these results indicated that intrinsic differences in the propensity for total particle production separate individuals, and that these are conserved across particle sizes.

**Figure 3.**
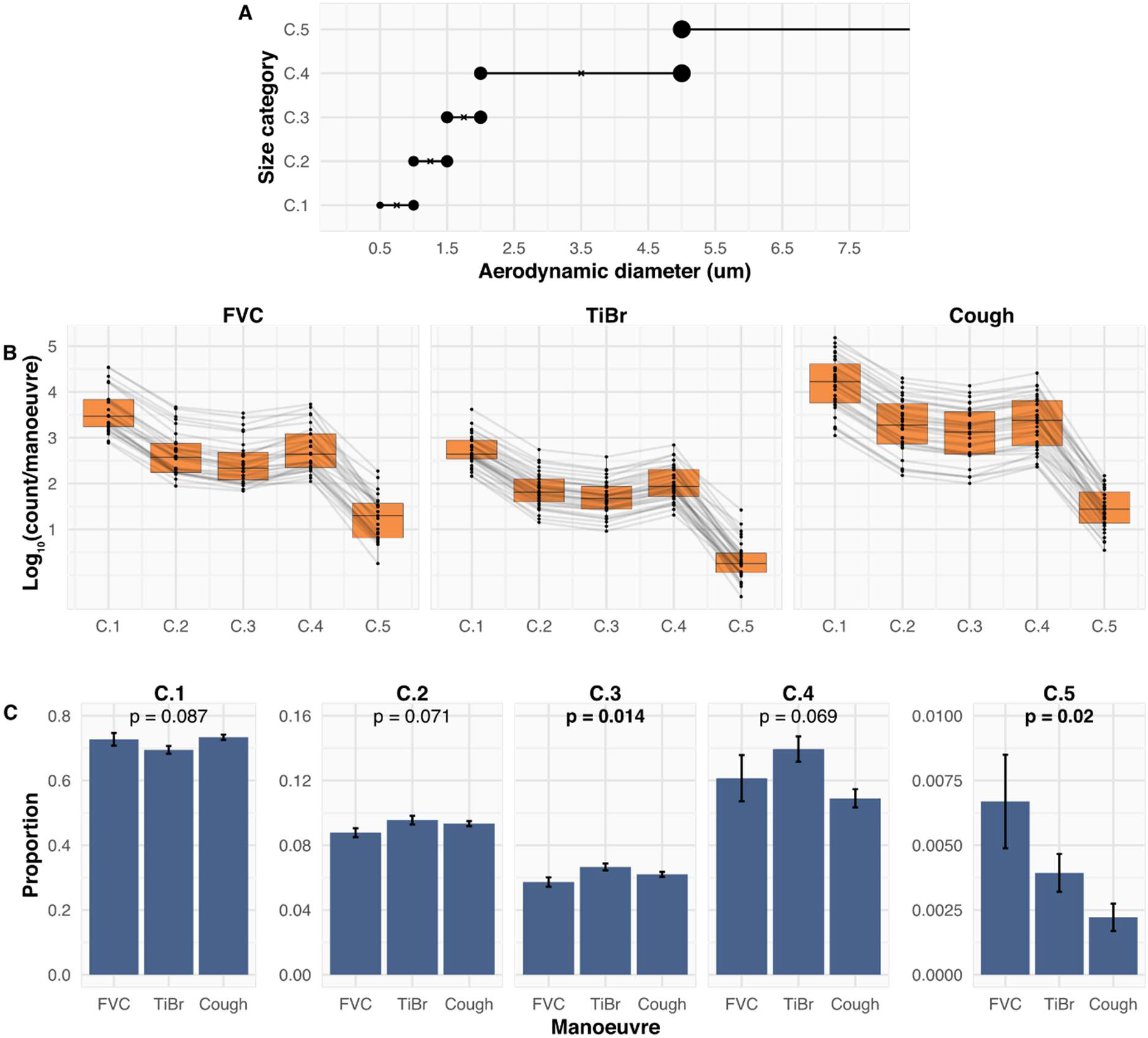
The relative contribution of particles of various sizes is consistent between FVC, TiBr and Cough. (**A**) Graphical representation of the size categories detected by the aerodynamic particle sizer. (**B**) A comparison of the average count per manoeuvre stratified by size category. Grey lines indicate the average number of particles per manoeuvre stratified by size category and participant ID (PTID). (**C**) A comparison between manoeuvres of the proportion composition of each size category per manoeuvre. The data in (**C**) are presented as the mean proportion ± SEM. A Repeated Measures ANOVA was performed, p values below 0.05 are represented in bold text.

### Use of three independent cyclone collectors to enable enumeration of Mtb bacilli

The relative contributions of different respiratory manoeuvres to aerosolization of *Mtb* bacilli has been poorly studied, with most reports focusing on Cough. We implemented a sampling strategy comprising 15 FVC and Cough manoeuvres and five minutes of TiBr. This resulted in closely matching volumes of bioaerosol collected for TiBr and FVC samples, with more than 3-fold the volume collected during Cough sampling (Figure E2C). Given the increased volume of bioaerosol collected from Cough and its assumed importance in TB transmission, we expected to find the greatest numbers of *Mtb* bacilli in the Cough samples.

The rate of production of *Mtb* was estimated for the three manoeuvres during the five minutes of sampling. For both FVC (IRR = 0.53, p = 0.0971) and Cough (IRR = 0.51, p = 0.0662), there was a trend to *Mtb* production at a lower rate compared to TiBr; however, neither of these was statistically significant (Figure 4A). Surprisingly, the percent of positive samples was consistent for all three manoeuvres, with 66%, 70%, and 65% of the samples positive for *Mtb* in TiBr, FVC, and Cough, respectively. Additionally, no significant differences were detected in the odds ratio of detecting *Mtb* between TiBr and FVC or TiBr and Cough (Figure 4B).

**Figure 4.**
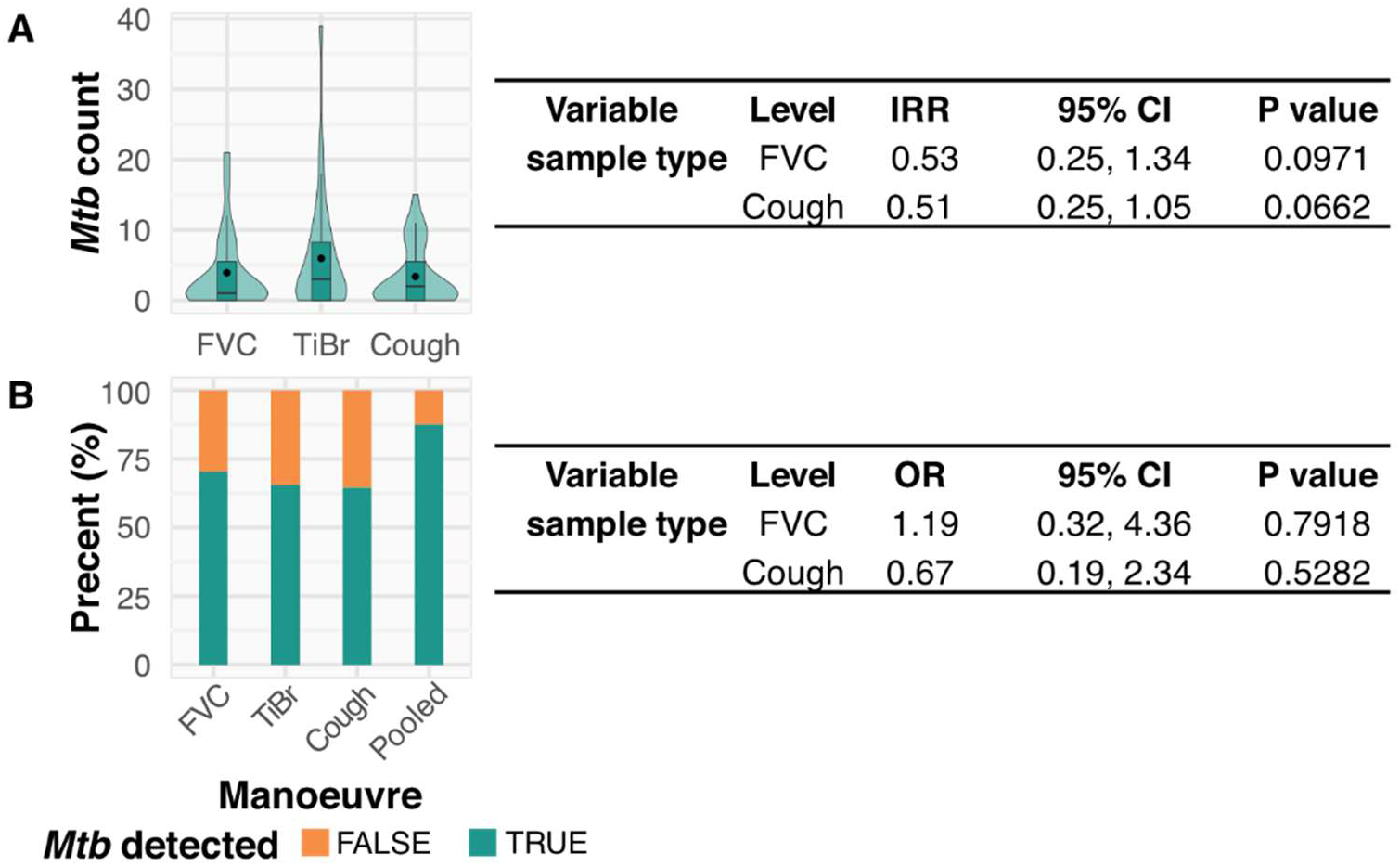
The detection of putative *Mycobacterium tuberculosis* within each respiratory manoeuvre sample. (**A**) A comparison of the total number of *Mycobacterium tuberculosis* (*Mtb*) detected between each sample, adjacent to the results from a negative binomial regression. (**B**) A comparison of the proportion of samples that were positive for aerosolised *Mtb*, adjacent to the results of a generalized linear mixed model. The “pooled” variable in (**B**) represents the percent of individuals that produced at least one positive sample.

The occurrence of spontaneous coughs during TiBr sampling might confound the TiBr measurement. To test this possibility, peaks detected in TiBr that were greater than 1.5 times the average peak height in the corresponding Cough sample were assumed to be spontaneous coughs (Figure E4A, right panel). Notably, plotting *Mtb* counts from TiBr against the number of coughs detected (Figure E4B) indicated no association between the number of *Mtb* and spontaneous coughs during sampling.

To examine the relationship between particle numbers and the aerosolization of *Mtb* bacilli, the relative abundance of *Mtb* bacilli per particle was calculated for all three manoeuvres. Participants with a zero count for *Mtb* were excluded from this analysis. The average concentration of *Mtb* bacilli for TiBr was 70% and 90% higher than that of FVC and Cough, respectively (Figure 5A). In addition, no correlation between total *Mtb* count and total particle count was observed for either FVC or Cough (Figure 5B). A slightly more apparent linear relationship was observed for TiBr (Figure 5B); however, this did not reach statistical significance. Together, these data imply a disconnect between the aerosolization of particles and *Mtb*.

**Figure 5.**
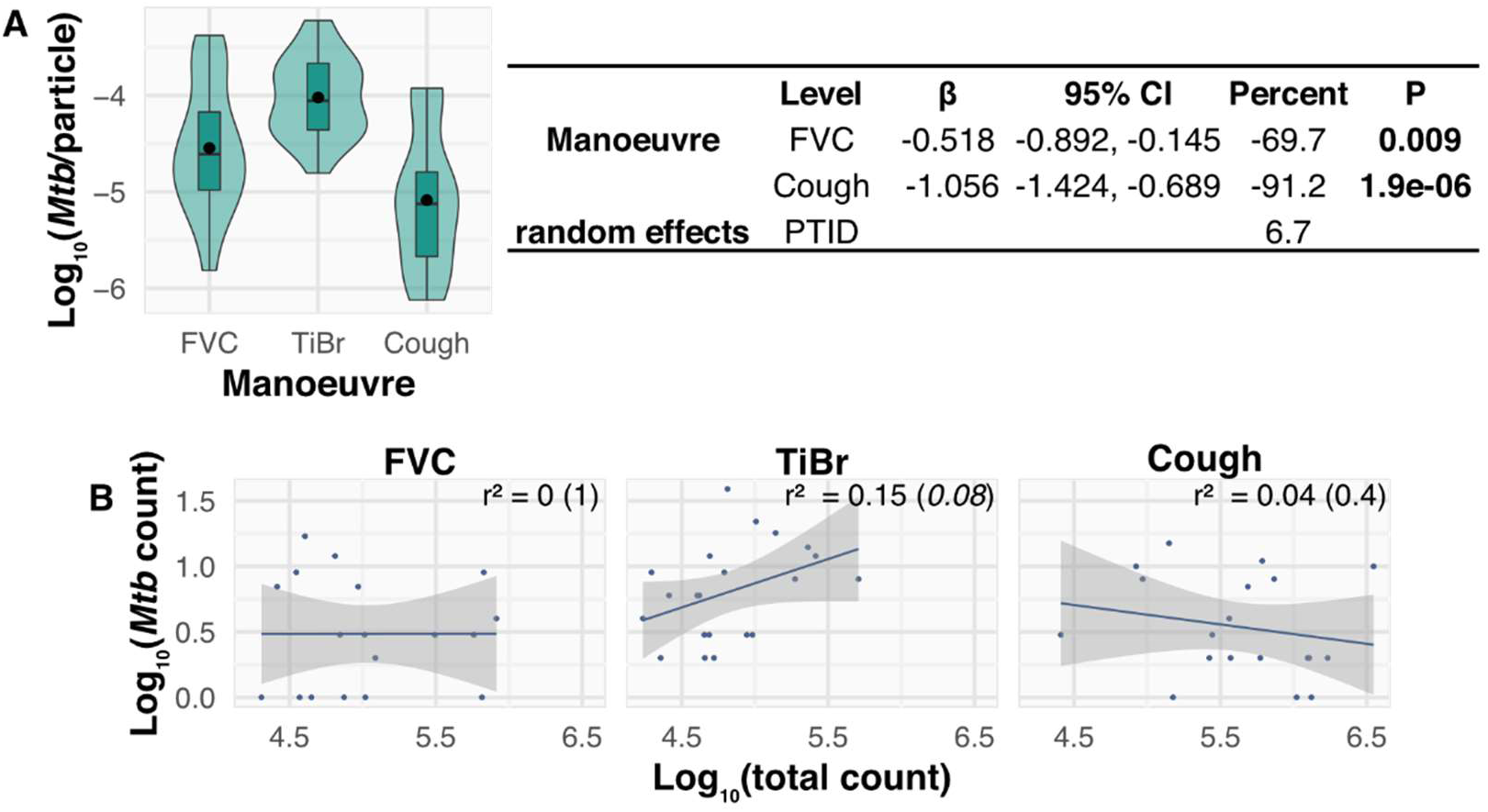
The concentration of *Mycobacterium tuberculosis* relative to particle count for each respiratory manoeuvre. (**A**) A comparison of the average number of *Mycobacterium tuberculosis* (*Mtb*) per particle within the bioaerosol, adjacent to the results from a linear mixed model. (**B**) Correlation assessment between log_10_(count) and log_10_(Mtb), with the results of a Pearson’s correlation (r-squared = r^2^ and p-value in brackets).

### The extent of aerosolization of Mtb depends predominantly on manoeuvre frequency

Knowing the concentration of *Mtb* per volume of bioaerosol suggested the potential to gain useful insight into the relative contributions of Cough and TiBr to the daily production of *Mtb*. To this end, we first compared the average number of bacilli produced per manoeuvre. On average, TiBr produced 2.6− and 3.2-fold fewer *Mtb* per manoeuvre compared to FVC and Cough, respectively (Figure 6A). Next, we extrapolated the values for the average number of bacilli per manoeuvre and the average frequency of manoeuvres per day to estimate daily *Mtb* production. Because FVCs are artificial, directed manoeuvres that are performed only under specific instruction, we utilized TiBr and Cough for this calculation.

**Figure 6.**
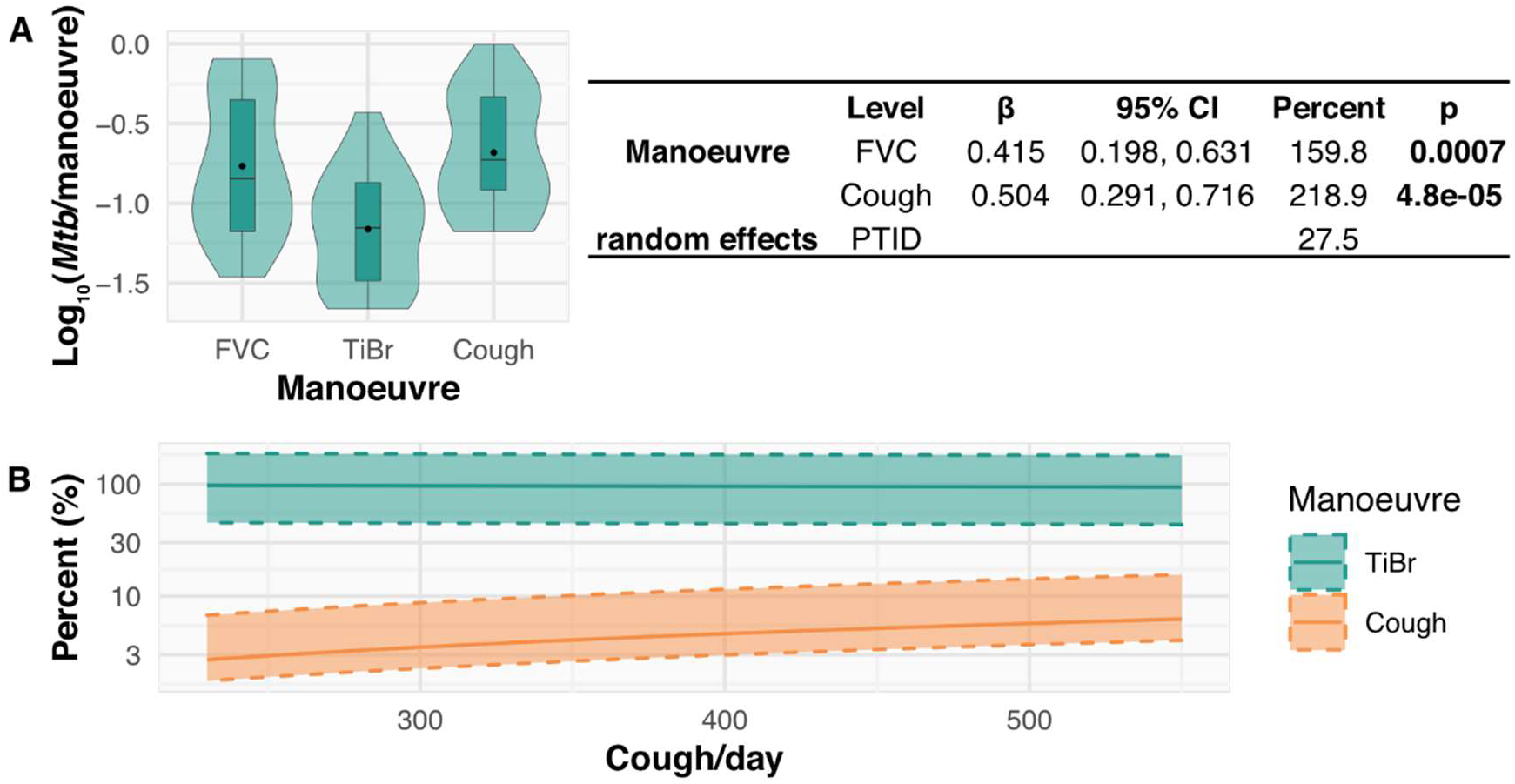
The relative contribution of *Mycobacterium tuberculosis* by each respiratory manoeuvre. (**A**) A plot of the average number of *Mycobacterium tuberculosis* (*Mtb*) per manoeuvre with the results of the linear mixed model. (**B**) The relative contribution of bacteria per day (percent). We used the median frequency of breaths to estimate an average of 22,047 breaths per day. We then assumed that for every Cough, there would be one fewer breath throughout the day over a range of coughs from 234 – 551 coughs. At each cough frequency, we determined the average number of bacilli per TiBr (solid green line) or per Cough (solid orange line). We then estimated the percentage contribution of aerosolised Mtb that each manoeuvre made relative to the total (solid lines sum to 100). The shaded regions represent the interquartile range of *Mtb* production for both TiBr and Cough.

Our data revealed that participant TiBr occurred at a rate of around one breath every 3.9 seconds, suggesting approximately 22,047 tidal breaths over 24 hours. Since we did not directly measure the frequency of spontaneous coughs, we estimated 24-hour Cough frequencies from published data (4), with a median of 466 coughs/day (first quartile 234, last quartile 551). We then used a constant maximum number of breaths per day (22,047) and assumed that, for each Cough, there would be one fewer breath. The average number of *Mtb* bacilli produced by TiBr and Cough was then calculated, and the relative proportion determined by dividing the number per manoeuvre by the total. This enabled an estimation of the relative contribution per day for an average person with an increasing number of coughs (Figure 6B).

It was apparent that Cough contributed between 3 and 7% of the total number of *Mtb* bacilli released, with TiBr consistently producing over 93%. Together, these data suggest that coughing is likely to produce significantly fewer bacilli per day compared to TiBr. That is, while the higher per event number and velocity of bacillary release might ensure an important role for coughing in disease transmission in short contacts, for typical exposures in high-risk settings such as public transport, workplaces, schools, *etc*. (13), TiBr is expected to contribute significantly to TB prevalence, especially in high-burden settings such as South Africa.

## Discussion

Cough has traditionally been considered the primary means of TB transmission (14). The result is that TB transmission research has predominately focused on factors including Cough production, frequency, and the Cough-borne *Mtb* bacillary load (2). However, the absence in all studies of a comparator respiratory manoeuvre (15–17) has rendered impossible any assessment of alternative contributory mechanisms. Transmission by aerosol requires the aerosolization of particles from the site of infection (18). For *Mtb*, which infects the peripheral lung and alveolar spaces (19), the proposed mechanism of particle aerosolization is fluid film rupture (20). According to this model, particles are produced during inspiration by alveolar reopening and released through expiration (21). Factors impacting particle release are therefore the rate of inspiration and the depth of expiration (21), with a recent study comparing deep exhalation and Cough finding no significant difference in the number of *Mtb* aerosolized between the two manoeuvres (10). For these reasons, we hypothesized that TiBr contributes to the aerosolization of *Mtb*. Therefore, we sought in this study to directly compare the propensity for particle and *Mtb* aerosolization via three defined respiratory manoeuvres: Cough, FVC and TiBr.

We sampled bioaerosols from 39 TB-positive participants. Consistent with findings from similar studies, 88% of participants produced at least one bioaerosol sample that was positive for *Mtb* (4, 8, 10), a marked increase over culture-based Cough-sampling techniques (17). Our results also indicated that all three respiratory manoeuvres were equally likely to produce *Mtb*, with TiBr, FVC, and Cough returning positive signals in 66%, 70%, and 65% of samples, respectively. When extrapolating based on daily manoeuvre frequency, these observations imply that TiBr contributes more than 90% of the daily aerosolised *Mtb* across a range of Cough frequencies – a conclusion consistent with the lack of correlation between *Mtb* aerosolization and Cough frequency (4).

Establishing a sampling algorithm appropriate for three distinct respiratory manoeuvres is challenging. However, the total number of particles produced during FVC and TiBr sampling were similar, with the Cough producing approximately 4-fold more particles. This suggested that, despite differences in sampling algorithms, the risk of under-sampling any manoeuvre was low. In addition, we saw significant variation between participants, spanning two orders of magnitude, consistent with previous observations (22). Per manoeuvre, TiBr produced significantly fewer particles than both FVC and Cough, with Cough producing the most particles. While it is tempting to speculate that the turbulence of the expired air played a role in the increased number of particles produced by Cough, this interpretation seems unlikely given the similarity in particle counts for Cough and FVC. Considering Cough and FVC are quite different in the rate of expiration, it might be more instructive that both these manoeuvres require deep inspiration: the inference, then, that the rate of expiration – and, therefore, the turbulence of expired air – plays a minimal role in aerosol generation is consistent with a fluid-film rupture model of aerosol generation in the peripheral lung (21). This is also consistent with the similarities in size distributions of particles between both participants and respiratory manoeuvres. Although the absolute counts per category varied between manoeuvres, the proportional compositions within each size category were conserved – an observation which supports the inference that the mechanism of particle production is consistent across the three respiratory manoeuvres (21, 22).

The average total number of *Mtb* per participant was 12.6 (max = 52), consistent with our previous study (8). However, owing to continued enhancements of our bioaerosol collection system, participants were sampled for ~15 minutes in this study *versus* the 60-minute sampling duration reported previously (8). Unexpectedly, all three respiratory manoeuvres produced consistently low levels of *Mtb*, with a mean count of 3.9 (max = 21), 5.9 (max = 39) and 3.4 (max = 15) for FVC, TiBr and Cough, respectively. TiBr samples tended to have a two-fold higher rate of *Mtb* aerosolization compared to both FVC and Cough; however, these differences were not significant. And, as noted above, the probability of a sample returning a positive result was consistent for all respiratory manoeuvres. Notably, among the participants who generated at least one positive sample, most (27/28) produced *Mtb* within their FVC and/or TiBr sample. These findings suggest that induced Cough may be unnecessary in studying *Mtb* transmission, a potentially important innovation given the strenuous nature of the induced Cough especially for unwell patients.

Contrary to our expectations, the concentration of *Mtb* bacilli per particle was 70% or 90% lower in FVC and Cough, respectively, compared to TiBr. In addition, no correlation was observed between particle number and *Mtb* count, even when stratified by participant. Together, these data suggest that variation in particle production alone is insufficient evidence to identify infectious patients, and that applications to reduce particle production seem unlikely to reduce infectiousness (23).

Despite the apparent unlinking of particle count and *Mtb* aerosolization, the sizeable increase in aerosol production during FVC and Cough manifest as a 3-fold increase in *Mtb* aerosolization for these manoeuvres compared to TiBr. However, when extrapolated to daily estimates, the relatively high frequency of breathing relative to coughing suggests that, over time, TiBr represents a major source of *Mtb* aerosols, as suggested previously (20). We calculated that, during any single day, breathing contributes >90% of the *Mtb* aerosolised by a TB-positive individual.

Our study had a number of limitations. The sample collection algorithm was not consistent for all respiratory manoeuvres – TiBr samples were primarily defined by time *versus* FVC and Cough that were defined by event number – and the particle collection and measurement apparatus were connected in parallel and not in series. Consequently, extrapolations were required to estimate the total number of particles and organisms present in the entire bioaerosol. Additionally, no work was done to determine the effect of manoeuvre order on particle or *Mtb* production. This could have impacted particle and *Mtb* production through participant exhaustion or through particle clearance. That said, we did separate FVC and Cough samples to ensure that TiBr provided a rest period between strenuous samples. The participants included in this study were diagnosed TB-positive via GeneXpert. Therefore, we cannot conclude the relative importance of TiBr to asymptomatic transmission. While our data indicate that significant levels of *Mtb* are aerosolised daily independent of participant Cough, further work is required to investigate this hypothesis in GeneXpert-negative, asymptomatic individuals. Owing to technical challenges inherent in studying spontaneous Cough over short sampling periods, we only studied induced Cough which may not be as infectious. Nevertheless, spontaneous coughs were detected in several TiBr samples, with no effect on overall production of *Mtb*. In estimating Cough frequency per hour, we assumed that the rate is consistent throughout the day. This is a strong assumption, and it is more likely that coughs cluster into discrete events with multiple coughs occurring at a time, suggesting potential outbursts of infectious aerosol production. A final limitation is that, while it might reasonably be assumed that *Mtb* bioaerosol counts are directly linked to infectiousness, this has not been formally demonstrated.

Despite these limitations, we interpret our results as indicating that TiBr might be a significant contributor to *Mtb* transmission in an endemic TB setting. This has significant ramifications for both transmission studies and intervention strategies. Firstly, bioaerosol sampling lends itself to a non-invasive participant sampling. Although the impact of induced Cough on a participant is relatively low, if a less invasive sampling algorithm can be applied, it should be. Secondly, interventions targeting disease transmission, such as active screening for symptomatic individuals, may not be effective. Therefore, linking bioaerosol organisms with infectious potential is of vital importance. Bioaerosol sampling is non-invasive and provides potential to identify infectious individuals well in advance of any typical screening regimen. This may offer a novel means to identify and treat infectious individuals before they manifest with definite symptoms.

A paradigm in which Cough is the primary driver of TB transmission places surpassing importance on lung pathology; moreover, it appears inconsistent with key epidemiological observations. Sub-clinical TB infections could represent a novel and uncontrolled source of disease transmission. Consequently, understanding how *Mtb* bacilli are aerosolised is of critical importance to curbing the epidemic in high-burden settings

## Author contribution

Conceptualization and design: RD, SG, AM, BL, JL, RS, DFW & RW; acquisition of data: JL, AM, BL, RS & RW; analysis and interpretation: RD, DFW & RW; first manuscript draft: RD & DFW; funding acquisition: DFW & RW. All authors critically reviewed and revised the manuscript for intellectual content and approved it prior to submission.

## Acknowledgements

We acknowledge the support of the South African Medical Research Council (SAMRC) with funds from National Treasury under its Economic Competitiveness and Support Package (MRC-RFA-UFSP-01-2013/CCAMP, RW), and the Strategic Health Innovations Partnerships (SHIP) Unit of the SAMRC (DFW) and as a sub-grant from the Bill and Melinda Gates Foundation (RW). We are grateful to the Bill & Melinda Gates Foundation (OPP1116641, RW), the US National Institutes of Health (NIH - R37AI058736, Freedberg PI; RW), the Research Council of Norway (R&D Project 261669 “Reversing antimicrobial resistance”, DFW), and the US National Institute of Child Health and Human Development (NICHD) U01HD085531 (DFW).

