## Supplementary material for "Aerosolization of *Mycobacterium tuberculosis* by tidal breathing": Online supplement

### *Online methods*

*Manoeuvre and peak detection*

$CO_2$ and particle data collection were contiguous throughout sampling (Figure 1A). Therefore, manoeuvre detection algorithms were written for each dataset to distinguish between FVC, TiBr and Cough sampling. A peak finding algorithm (E1) was run on relative $CO_2$ and particle counts, with a bespoke local smoothing algorithm applied to smooth peaks. Peaks (individual manoeuvres) were then checked visually. In 12 participants, peaks were manually adjusted. For FVC and Cough samples, peak identification was critical, as only particles that fell within these peaks were considered. This was because participants removed their heads from the sampling apparatus between manoeuvres. For TiBr, the participants' heads remained in the cone for the duration of sampling; therefore, all particles were considered for TiBr samples.

*Volume calculation for bioaerosol particles*

The APS enumerated particles in five size categories, designated C.1 to C.5. The diameter ranges for these categories were 0.5 – 1 µm (C.1), 1 – 1.5 µm (C.2), 1.5 – 2 µm (C.3), 2 – 5 µm (C.4) and >5 µm (C.5) (Figure 3A). Particles were assumed to be spherical and the mean diameter from each size category was estimated without curve fitting. For C.1, this was assumed to be 0.5 µm as counts in this category were on average 0.892 $\log_{10}$-units higher than in C.2, suggesting a greater proportion of smaller particles. The differences in C.2 – C.4 were smaller; therefore, the mid-point of each bin was used to estimate average diameter. A diameter of 5 µm was used for C.5. Bioaerosol volume was calculated with the following formula:

$$bioaerosol\ volume\ (nL)_{C.x} = \frac{\frac{4}{3} \times \pi \times \left(\frac{diameter_{C.x}}{2}\right)^3 \times\ count_{C.x}}{10^6}$$

*Estimation of per manoeuvre particle counts and volumes*

The flow rate via the aerosol particle sizer (APS) was constant for all manoeuvres and only differed by duration (Figure 1A). For TiBr, the flow rate via the APS was one-third that of the total bioaerosol sample. Therefore, to estimate the total number of particles released during TiBr, the count for each sample was multiplied by three. This gave the overall sample count for each manoeuvre (Figure 1B). The sample count was divided by the number of manoeuvres detected within the particle data to determine the average number of particles per manoeuvre (count/manoeuvre) for each sample. The count/manoeuvre variable was then multiplied by the total number of peaks detected in the $CO_2$ data for each participant to obtain the estimated total count for both FVC and Cough samples. This led to a better comparison of the total number of particles produced across the three manoeuvres.

*Estimation of the total number of Mtb bacilli*

For TiBr, only two-thirds of the bioaerosol was collected *versus* eight-ninths for both FVC and Cough. To estimate the total number of *Mtb* bacilli in each sample, the microscope counts were multiplied by 3/2 for TiBr and 9/8 for FVC and Cough.

*Linear Mixed Models*

Individual participants each produced bioaerosol samples from three respiratory manoeuvres, violating the assumption of independence and necessitating an alternative approach. Various linear mixed-effects models were applied as the addition of the random effect for slope enabled the average difference between manoeuvres to be determined while accounting for variation between participants. All linear mixed models outlined below contained manoeuvre (sample type) as the “fixed effect” and participant ID (PTID) as the “random effect”. This was done using the lme4 package in R (E2). Where indicated, additional fixed effects were added.

*Linear mixed effects models*

To determine manoeuvre differences in average particle count, volume, *Mtb*/volume and *Mtb*/manoeuvre, these outcome variables were $\log_{10}$-transformed and linearity, normality of residuals, and homoskedasticity assessed. Next, they were regressed against sample type in separate univariate regression models. The simplified equation for a simple linear regression is:

$$y = \beta_0 + \beta_1 X_1$$

However, owing to the outcome variable being $\log_{10}$-transformed, the equation was modified to:

$$\log_{10}(y) = \beta_0 + \beta_1 X_1$$

In the raw form, the β coefficients are therefore interpretated as a unit change in $X_1$ leading to a $\beta_1$ change in $\log_{10}(y)$. Alternatively, the coefficients can be modified:

$$y = 10^{\beta_0} + 10^{\beta_1 X_1}$$

This provides a slightly more intuitive interpretation whereby a unit change in $\beta_1$ relates to a fold change in y, relative to TiBr. The following equation provides the percentage change in y relative to TiBr:

$$Percentage\ change = \left(10^{\beta 1} - 1\right) \times 100$$

*Generalized linear mixed models*

For the binary outcome of sample positivity for putative *Mtb*, logistic regression was performed with sample type as the fixed effect and variation in slope (random effects) accounted for with PTID. The odds ratio (OR) was determined by exponentiating the $\beta_1$ coefficient:

$$OR = e^{\beta 1}$$

*Negative binomial regression*

For count data obtained through enumerating *Mtb* in each sample, a negative binomial regression was applied using the MASS package in R (E3). Owing to the complexity of incorporating random effects into this model and the poor correlation between individual and *Mtb* count across the different manoeuvres, random effects were not considered, and statistical independence was assumed. As overall sampling duration was set to about five minutes, no offset was applied for this analysis. As such, this regression model simply interrogated whether there were different rates of *Mtb* production within the different samples, not accounting for other potential differences. Exponentiating the $\beta_1$ coefficient gave the incident rate ratio (IRR):

$$IRR = e^{\beta 1}$$

Assessment of model appropriateness was tested with the deviance goodness of fit test, with a visual inspection done by plotting the mean and dispersion parameter calculated within the model overlayed on the data.

*Correlation analysis*

To assess the correlation between putative *Mtb* count and bioaerosol count, a correlation analysis was conducted with a $\log_{10}$-transformation of both variables. Negative counts (zero *Mtb* detected) were excluded. Linearity was visually assessed, and a Pearson correlation was performed.

### *Online methods references*

E1. Borchers HW. pracma: Practical Numerical Math Functions. R package version 2.3.3; 2021.

E2. Douglas B, Martin M, Ben B, Steve W. Fitting Linear Mixed-Effects Models Using lme4. *Journal of Statistical Software* 2015; 67: 1-48.

E3. Venables WN, Ripley BD. Modern Applied Statistics with S. New York; 2002.

### *Supplemental figures*

### *Figure E1*

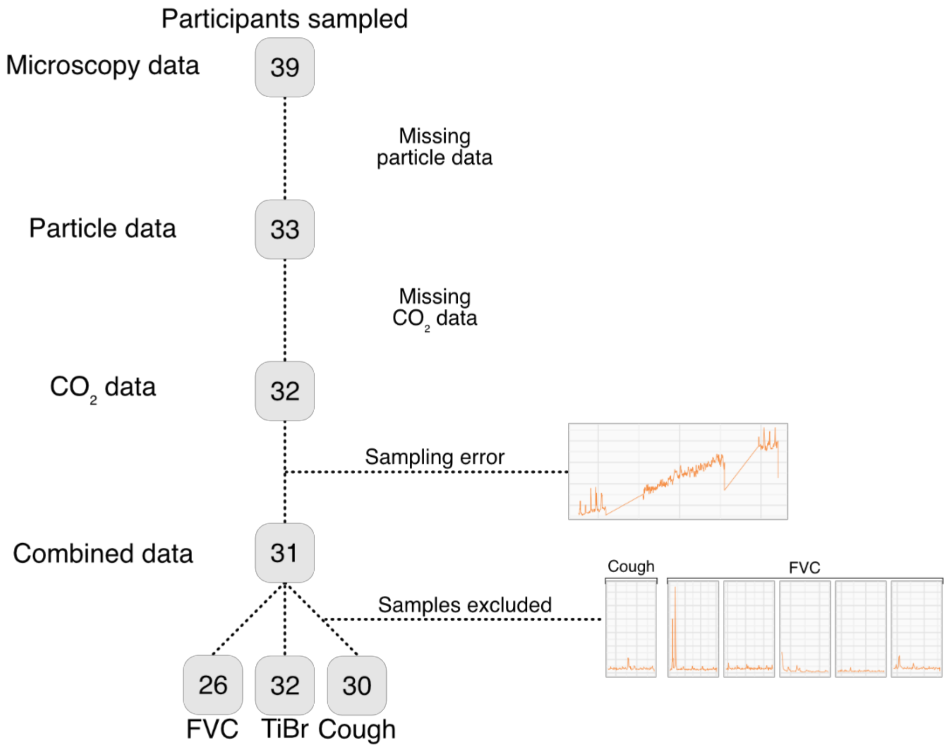

**Figure E1. Participants and samples that were excluded.**

### *Figure E2*

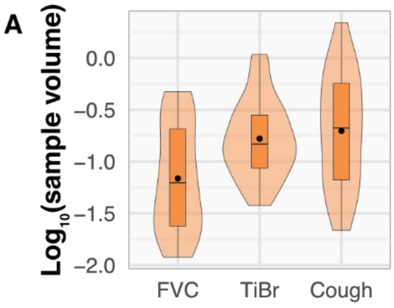

| | Level | β | 95% CI | Percent | p |
|---|---|---|---|---|---|
| **Manoeuvre** | FVC | -0.387 | -0.615, -0.158 | -59 | **0.0015** |
| | Cough | 0.075 | -0.144, 0.294 | 19 | 0.5039 |
| **random effects** | PTID | | | 15.3 | |

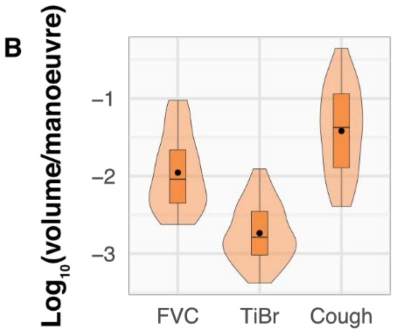

| | Level | β | 95% CI | Percent | p |
|---|---|---|---|---|---|
| **Manoeuvre** | FVC | 0.777 | 0.551, 1.004 | 498.9 | **6.5e-09** |
| | Cough | 1.317 | 1.100, 1.534 | 1975 | **<2e-16** |
| **random effects** | PTID | | | 15.7 | |

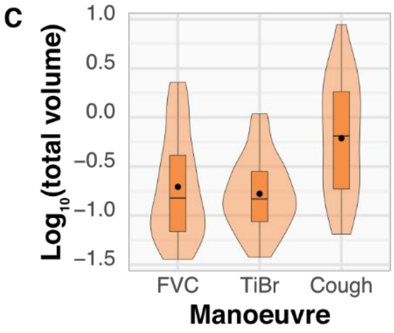

| | Level | β | 95% CI | Percent | p |
|---|---|---|---|---|---|
| **Manoeuvre** | FVC | 0.070 | -0.169, 0.308 | 17.4 | 0.569 |
| | Cough | 0.568 | 0.340, 0.797 | 270.0 | **8.8e-06** |
| **random effects** | PTID | | | 16.5 | |

**Figure E2. Variation in particle production by FVC, TiBr and Cough.** A comparison of the (**A**) sample volume, (**B**) volume/manoeuvre and (**C**) total volume of particles during sampling. The adjacent tables contain the results of univariate linear mixed models for each. The beta-coefficient (β) and 95% confidence interval (CI) are presented with percentage change relative to TiBr (Percent). The random effects results indicate the degree of variation (in %) between participants.

### *Figure E3*

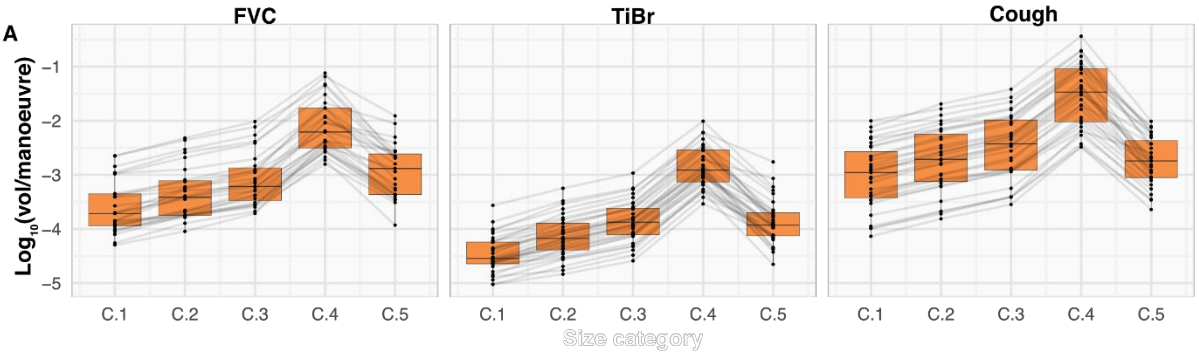

**B**

| | Level | β | 95% CI | Percent |
|---|---|---|---|---|
| **Manoeuvre** | FVC | **0.820** | 0.727, 0.913 | 560.5 |
| | Cough | **1.338** | 1.250, 1.427 | 2079.5 |
| **Size category** | C2 | **-0.892** | -1.009, -0.776 | -83.2 |
| | C3 | **-1.069** | -1.186, -0.952 | -88.8 |
| | C4 | **-0.798** | -0.915, -0.682 | -79.2 |
| | C5 | **-2.489** | -2.606, -2.372 | -99.6 |
| **random effects** | PTID | | | 31.3 |

**C**

| | Level | β | 95% CI | Percent |
|---|---|---|---|---|
| **Manoeuvre** | FVC | **0.820** | 0.727, 0.913 | 560.5 |
| | Cough | **1.338** | 1.250, 1.427 | 2079.5 |
| **Size category** | C2 | **0.302*** | 0.185, 0.418 | 100.2 |
| | C3 | **0.563** | 0.447, 0.680 | 265.9 |
| | C4 | **1.536** | 1.420, 1.653 | 3337.8 |
| | C5 | **0.511** | 0.394, 0.628 | 224.3 |
| **random effects** | PTID | | | 31.3 |

**Figure. E3. The relative volume contribution of particles of various sizes is consistent between FVC, TiBr and Cough.** (**A**) A comparison of the average volume per manoeuvre stratified by size category. Grey lines indicate the average volume of particles per manoeuvre stratified by size category and participant ID (PTID). Results for a mixed effects linear regression of average count (**B**) or volume (**C**) per manoeuvre against size category and manoeuvre. P < 2e-16 for all coefficients, except where indicated by an asterisk, p = 5.63e-07.

### *Figure E4*

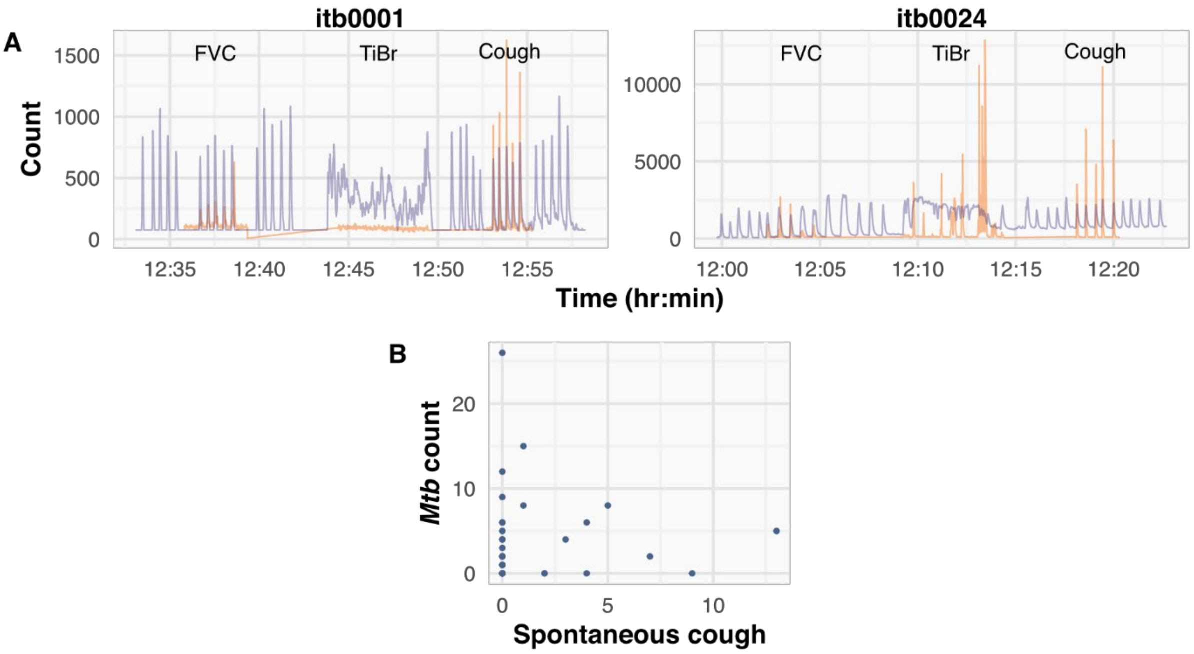

**Figure E4. No association was detected between spontaneous coughs during TiBr and the release of aerosolised *M. tuberculosis*.** (**A**) $CO_2$ (purple) and particle count (orange) *vs* time stratified by participant. Itb0001 represents an ideal sample collection. Itb0024 indicates detection of spontaneous coughing during TiBr sampling. (**B**) Assessment of relationship between putative *Mtb* and the detection of coughing within TiBr samples
